# Potassium channels contribute to activity-dependent scaling of dendritic inhibition

**DOI:** 10.1101/098889

**Authors:** Jeremy T. Chang, Michael J. Higley

## Abstract

GABAergic inhibition plays a critical role in the regulation of neuronal activity. In the neocortex, inhibitory interneurons that target the dendrites of pyramidal cells influence both electrical and biochemical postsynaptic signaling. Voltage-gated ion channels strongly shape dendritic excitability and the integration of excitatory inputs, but their contribution to GABAergic signaling is less well understood. By combining 2-photon calcium imaging and focal GABA uncaging, we show that voltage-gated potassium channels normally suppress the GABAergic inhibition of calcium signals evoked by back-propagating action potentials in dendritic spines and shafts of cortical pyramidal neurons. Moreover, the voltage-dependent inactivation of these channels leads to enhancement of dendritic calcium inhibition following somatic spiking. Computational modeling reveals that the enhancement of calcium inhibition involves an increase in action potential depolarization coupled with the nonlinear relationship between membrane voltage and calcium channel activation. Overall, our findings highlight the interaction between intrinsic and synaptic properties and reveal a novel mechanism for the activity-dependent scaling of GABAergic inhibition.

**Significance Statement:** GABAergic inhibition potently regulates neuronal activity in the neocortex. How such inhibition interacts with the intrinsic electrophysiological properties of single neurons is not well-understood. Here we investigate the ability of voltage-gated potassium channels to regulate the impact of GABAergic inhibition in the dendrites of neocortical pyramidal neurons. Our results show that potassium channels normally reduce inhibition directed towards pyramidal neuron dendrites. However, these channels are inactivated by strong neuronal activity, leading to an enhancement of GABAergic potency and limiting the corresponding influx of dendritic calcium. Our findings illustrate a previously unappreciated relationship between neuronal excitability and GABAergic inhibition.

## Introduction

Inhibition in the neocortex is primarily mediated by the neurotransmitter gamma-aminobutyric acid (GABA) through synaptic contacts made by interneurons. These synapses are distributed across the entire somatodendritic arbor and work to counteract excitatory glutamatergic input. GABAergic synapses that target the axon initial segment and soma exert a strong influence on somatic voltage, and consequently play important roles in regulating the generation and timing of action potentials (1–4). However, the vast majority of inhibitory inputs are formed onto pyramidal cell dendrites (5), and the role of dendrite-targeting inhibition has been an area of growing interest (6–8).

One important function of dendritic inhibition, in addition to action potential regulation, is the regulation of dendritic calcium signals which are thought to play an instructive role in synaptic plasticity (9, 11). Recent reports in the neocortex and hippocampus have described varying efficacy of dendritic calcium inhibition, ranging from spatial compartmentalization within individual spines to complete abolition of actively back-propagating action potentials (bAPs)(10, 12–15). The mechanisms underlying the heterogeneity of previous findings are unclear, but one contributing factor may be variation in intrinsic dendritic properties, like voltage-dependent channels, whose impact on GABAergic inhibition of bAPs is not well understood. Indeed, earlier work in hippocampal neurons suggested that the inhibition of bAP-evoked dendritic calcium signaling may be inversely correlated with the magnitude of the calcium transient (14), consistent with an interaction between GABAergic potency and dendritic excitability.

The expression of voltage-gated ion channels within neuronal dendrites regulates cellular excitability and strongly influences synaptic integration (16–20). Potassium channels, including those sensitive to the blocker 4-aminopyridine (4-AP), are expressed throughout the dendritic arbors of cortical and hippocampal pyramidal neurons and have been implicated in the regulation of back-propagating action potentials (bAPs) and excitatory synaptic integration (21–26). Interestingly, A-type Kv4.2 channels have been shown to preferentially co-localize with GABAergic synapses, suggesting they may also play a role in the control of inhibition (27, 28).

Here, we examine how voltage-gated potassium channels alter GABAergic inhibition of bAP-evoked Ca2+ signals ΔCa2+) in dendrites of L5 pyramidal neurons in mouse visual cortex. We show that the blockade of these channels enhances both the amplitude of bAP-evoked ΔCa2+ and unexpectedly also the inhibition of bAP-evoked ΔCa2+. We also show that the voltage-dependent inactivation of these channels gives rise to a scaling of dendritic GABAergic inhibition, such that inhibitory efficacy is enhanced following strong somatic activity. Thus, our findings demonstrate that intrinsic excitability interacts with GABAergic synaptic input to dynamically regulate dendritic Ca2+ signaling.

## Results

In order to investigate the impact of potassium channels on dendritic inhibition, we performed two-photon calcium imaging of bAP-evoked dendritic Ca2+ transients in layer 5 pyramidal neurons (L5PNs) of mouse visual cortex (Figure 1A). Ca2+ signals were measured in dendritic spines and neighboring shafts along the primary apical dendrite, 100-150 μη from the soma (Figure 1B). To probe the effects of GABAergic inhibition on ΔCa2+, we compared uninhibited ΔCa2+ from bAPs induced by somatic current injection with ΔCa2+ from bAPs preceded (15 ms) by local (at the imaging site) uncaging of RuBi-GABA (29). To compare observations across different recordings, GABAergic inhibition of ΔCa2+ was quantified as in previous studies as ΔCa2+CtlΔCa2+Inh)MCa2+al (10). The magnitude of this Ca2+ inhibition was measured before and after bath-application of the potassium channel blocker 4-aminopyridine (4-AP)(Figure 1A). Treatment with 4-AP broadened the somatic action potential (p=0.0020)(Figure 1C) and increased the average peak ΔCa2+ evoked by a single bAP for both spines (p=0.0059) and neighboring shafts (p=0.0137) (Figure 1D-G). Moreover, 4-AP significantly increased the amplitude of the uncaging-evoked inhibitory postsynaptic potential (IPSP, Fig. 1C, p=0.0391) and enhanced the average GABAergic inhibition of ΔCa2+ for both spines (p=0.0195) and neighboring shafts (p=0.0098) (Figure 1D-G). This result was not observed when slices were pre-treated with the GABAaR antagonist picrotoxin (data not shown).

**Figure 1.**
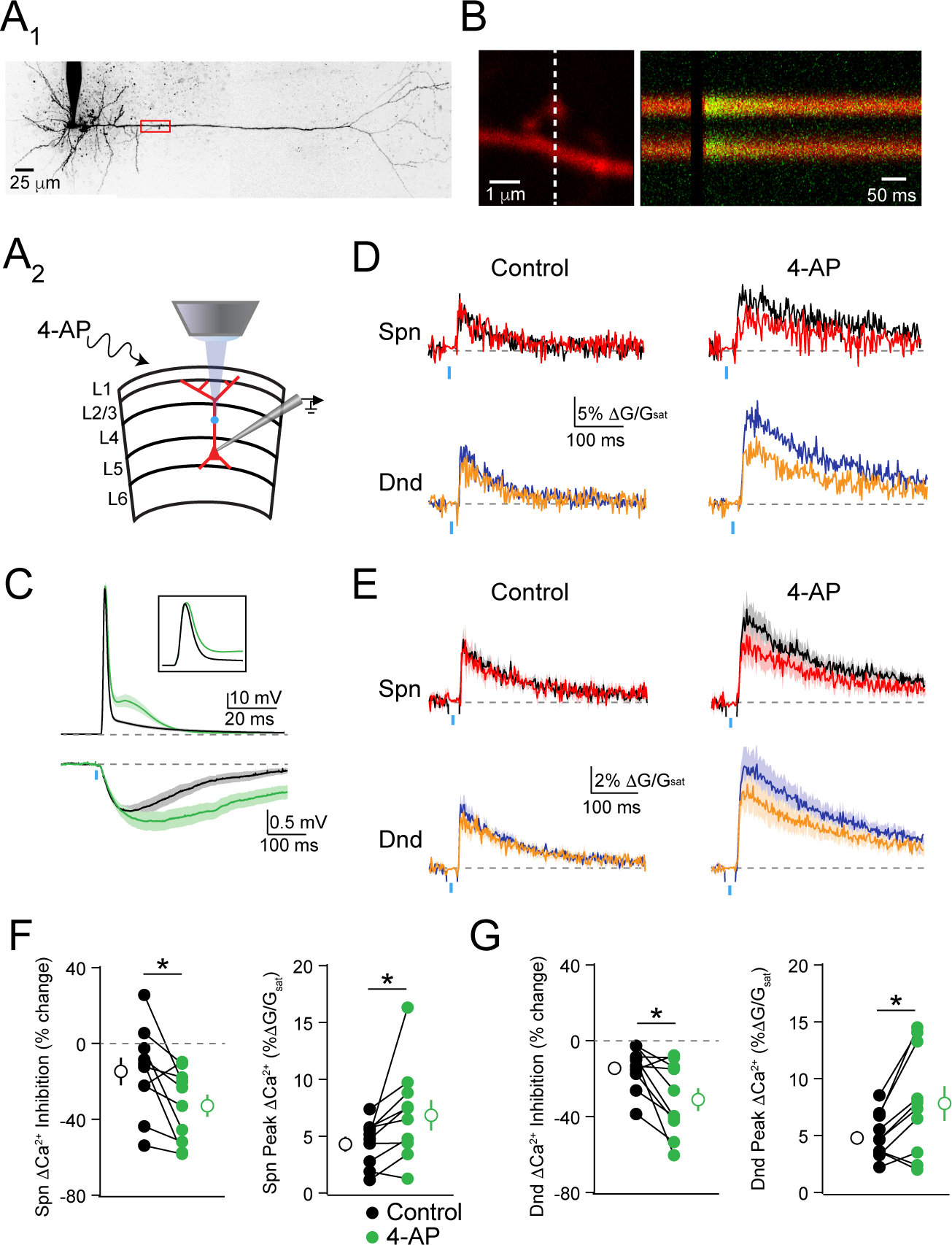
GABAergic inhibition of Δ Ca2+ is enhanced by blockade of potassium channels. (**A**) Whole Cell patch recordings were performed in L5PNs of visual cortex. Ca2+ imaging and GABA uncaging were performed along the proximal apical dendrite, as shown in the example cell (A1) and schematic (A2). (**B**) Example spine-dendrite pair and the associated line-scanned response to a bAP. (C) Average ± SEM somatic voltage recorded before (black) and after (green) treatment with 4-AP for action potentials (upper traces) and uncaging-evoked IPSPs (lower traces). (D) Example bAP-evoked ΔCa2+ for the apical dendritic region shown in (**B**) for bAP alone (black, blue) or paired with GABA uncaging (red, orange) for the spine and neighboring dendrite, before (left) and after (right) treatment with 5 mM 4-AP. (E) Average (n=10) ΔCa2+ for the population of imaged spines and dendrites, colors as in (D). (F-G) Population data (n=10) showing the magnitude of ΔCa2+ inhibition and peak bAP-evoked ΔCa2+ for spines (F) and neighboring dendrites (G) before (black) and after (green) treatment with 5 mM 4-AP (Wilcoxon matched-pairs signed rank test, *p<0.05).

Next, we investigated whether the actions of 4-AP on dendritic Ca2+ inhibition required block of potassium channels near the site of GABAergic input. We used a puffer pipette to locally apply 4-AP at different locations along the somatodendritic axis. When applied to the proximal apical dendrite (at the site of GABA uncaging), 4-AP replicated the effects of bath-application on the magnitude of bAP-evoked ΔCa2+ and its inhibition by GABA. Specifically, 4-AP increased ΔCa2+ in spines (p=0.0156) and neighboring shafts (p=0.0313) (Figure 2A-D). GABAergic inhibition of ΔCa2+ was also enhanced in spines (p=0.0469) and neighboring shafts (p=0.0313) (Figure 2A-D). In contrast, application of 4-AP to the cell body had no impact on peak ΔCa2+ or its inhibition by GABA within the proximal apical dendrite (Supplemental Fig. 1). Thus, our data suggest that the impact of 4-AP on GABAergic inhibition of Ca2+ is mediated by a distinct pool of dendritic potassium channels.

**Figure 2.**
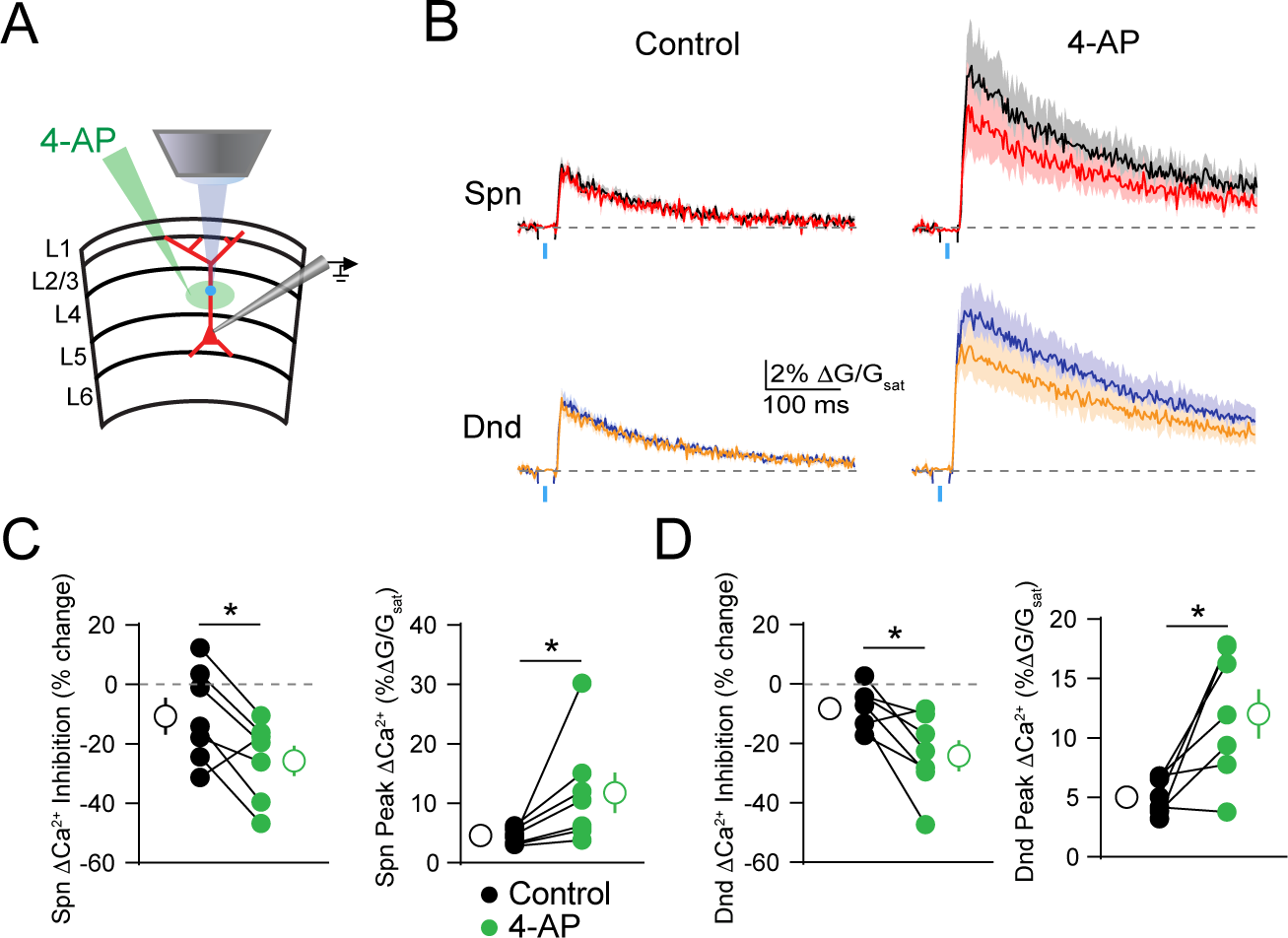
GABAergic inhibition of Δ Ca2+ is enhanced by local blockade of potassium channels. (**A**) Schematic of recording and imaging configuration, illustrating location of puffed 4-AP. (**B**) Average ± SEM (n=7) Ca2+ transients for bAP alone or paired with GABA uncaging before (left) and after (right) application of 4-AP to the proximal apical dendrite (colors as in Fig. 1). (**C-D**) Population data (n=7) showing the magnitude of Ca2+ inhibition and peak bAP-evoked Ca2+ for spines (C) and neighboring dendritic shafts (D) in control (black) and 4-AP (green) conditions (Wilcoxon matched-pairs signed rank test, *p<0.05).

One feature of many potassium channels is their voltage-dependent inactivation, which limits their conductance during periods of high neuronal activity (30, 31). We therefore asked whether this property might enable dendritic GABAergic inhibition to dynamically scale with somatic firing. To test this hypothesis, we compared GABAergic inhibition of ΔCa2+ evoked by a bAP alone or preceded 20 ms by a train of 5 bAPs at 100 Hz (Figure 3A). Similar to 4-AP, the preceding train significantly enhanced the peak ΔCa2+ for both spines (p=0.002) and neighboring shafts (p=0.002) and also enhanced GABAergic inhibition of ΔCa2+ for spines (p=0.0059) and neighboring shafts (p=0.002)(Figure 3B-D). Importantly, the ability of the 100 Hz train to enhance dendritic inhibition was occluded by prior bath application of 4-AP (Supplemental Fig. 2), suggesting that both manipulations target a similar population of dendritic potassium channels.

**Figure 3.**
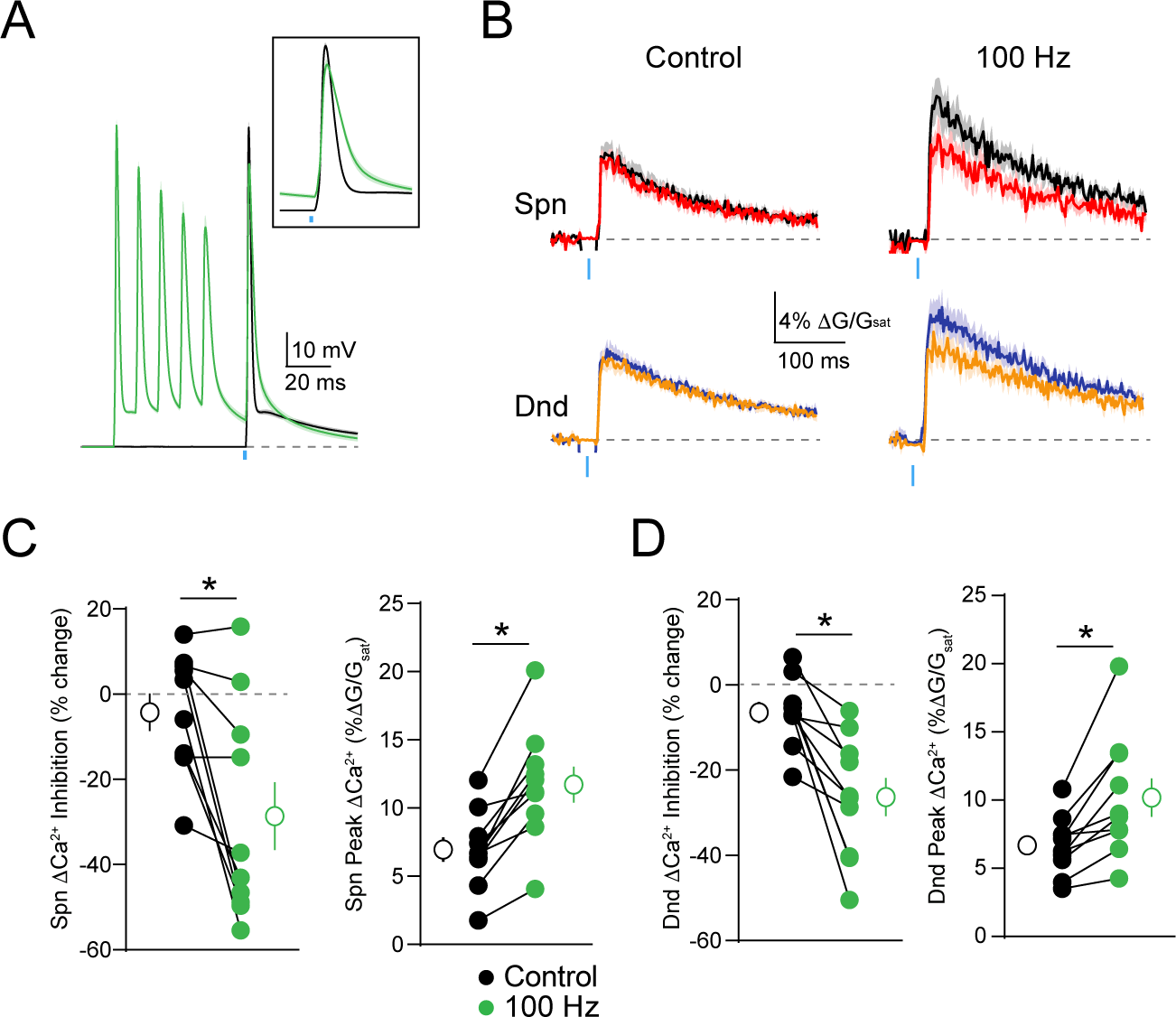
Somatic activity enhances GABAergic inhibition of dendritic Δ Ca2+. (**A**) Average ± SEM (n=10) of somatic recordings for a single action potential (black) or when preceded 20 ms by a 100 Hz train of action potentials (green). Inset shows change in spike waveform. (**B**) Average ± SEM (n=10) Ca2+ transients for bAP alone or paired with GABA uncaging, presented either singly (left) or following a 100 Hz train of action potentials (right)(colors as in Fig. 1). (**C-D**) Population data (n=10) showing the magnitude of Ca2+ inhibition and peak bAP-evoked Ca2+ for spines (C) and neighboring dendritic shafts (D), presented either singly (black) or following a 100 Hz train of action potentials (green)(Wilcoxon matched-pairs signed rank test, *p<0.05).

We next asked whether activity-dependent scaling of inhibition could be seen with synaptic GABA release. To test this, we expressed channelrhodopsin-2 (ChR2) in a subset of dendrite-targeting cortical interneurons expressing somatostatin (SOM-INs)(Figure 4A). Brief pulses of blue light were used to activate SOM-INs and produce postsynaptic IPSPs. We repeated experiments comparing the GABAergic inhibition of ΔCa2+ evoked by a bAP alone or preceded by a 100 Hz train. As with GABA uncaging, trains of somatic action potentials significantly enhanced the magnitude of ΔCa2+ in spines (p=0.0098) and neighboring shafts p=0.0059) and led to stronger GABAergic inhibition of ΔCa2+ for both spines (p=0.0273) and neighboring shafts (p=0.0273) (Figure 4B-D). Taken together, these results demonstrate that voltage-dependent potassium channels in the apical dendrite play a key role in shaping the impact of synaptic GABAergic inhibition on bAP-evoked ΔCa2+.

**Figure 4.**
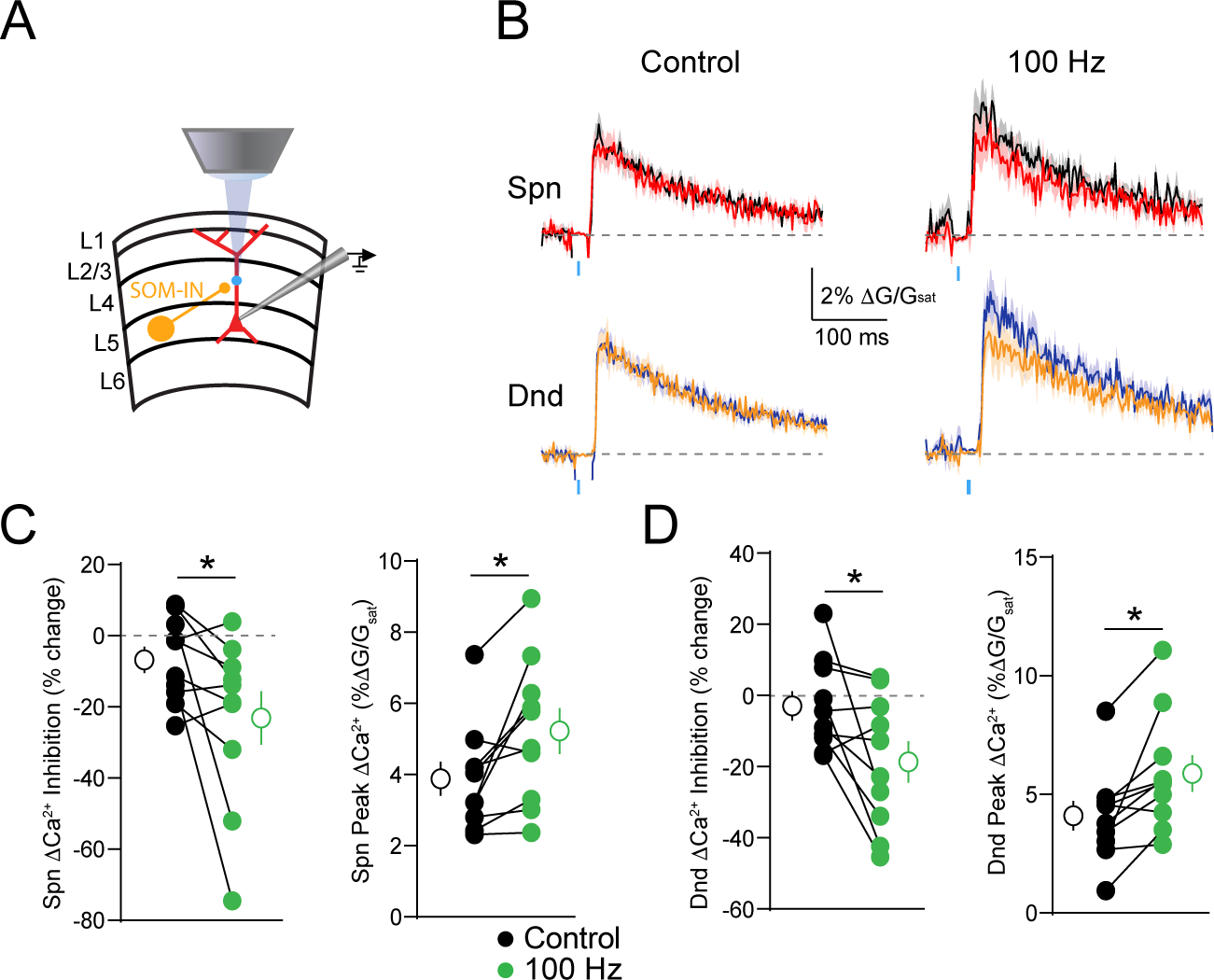
Somatic activity enhances synaptic GABAergic inhibition of Δ Ca2+. (**A**) Schematic showing recording and imaging configuration. ChR2 was virally expressed in somatostatin-containing interneurons (SOM-INs) and activated with blue light pulses. (**B**) Average ± SEM (n=10) Ca2+ transients for bAP alone or paired with optical stimulation of SOM-INs, presented either singly (left) or following a 100 Hz train of action potentials (right)(colors as in Fig. 1). (**C-D**) Population data (n=10) showing the magnitude of Ca2+ inhibition and peak bAP-evoked Ca2+ for spines (C) and neighboring dendritic shafts (D), presented either singly (black) or following a 100 Hz train of action potentials (green)(Wilcoxon matched-pairs signed rank test, *p<0.05).

Finally, to examine the biophysical mechanisms underlying the interaction of dendritic potassium channels with GABAergic signaling, we simulated an active dendritic compartment (see Methods) and tested the impact of varying an A-type potassium conductance (gK_A_) on the magnitude of Ca2+ inhibition (Figure 5). We found that varying the maximal dendritic gK_A_ betw een control (70 mS/cm2 in the distal compartment, 17.0 mS/cm2 at the synapse) and low (10 mS/cm2 in the distal compartment, 2.4 mS/cm2 at the synapse) conditions, thus simulating application of 4-AP, recapitulated our experimental data, increasing the peak AP amplitude and enhancing Ca2+ inhibition. (Fig. 5A). Interestingly, the reduction in peak AP amplitude caused by GABAergic inhibition was not strongly affected by reducing gK_A_ (Fig. 5B). However, we found that the relationship between peak AP amplitude and peak Ca2+ current was highly supralinear (Fig. 5C). Thus, a similar amount of GABAergic shift in the peak membrane potential produced a substantially larger inhibition of Ca2+ influx when gK_A_ was reduced (Fig. 5C). Our model therefore demonstrates a straightforward biophysical mechanism linking potassium conductance, GABAergic signaling, and Ca2+ inhibition in PN dendrites.

**Figure 5.**
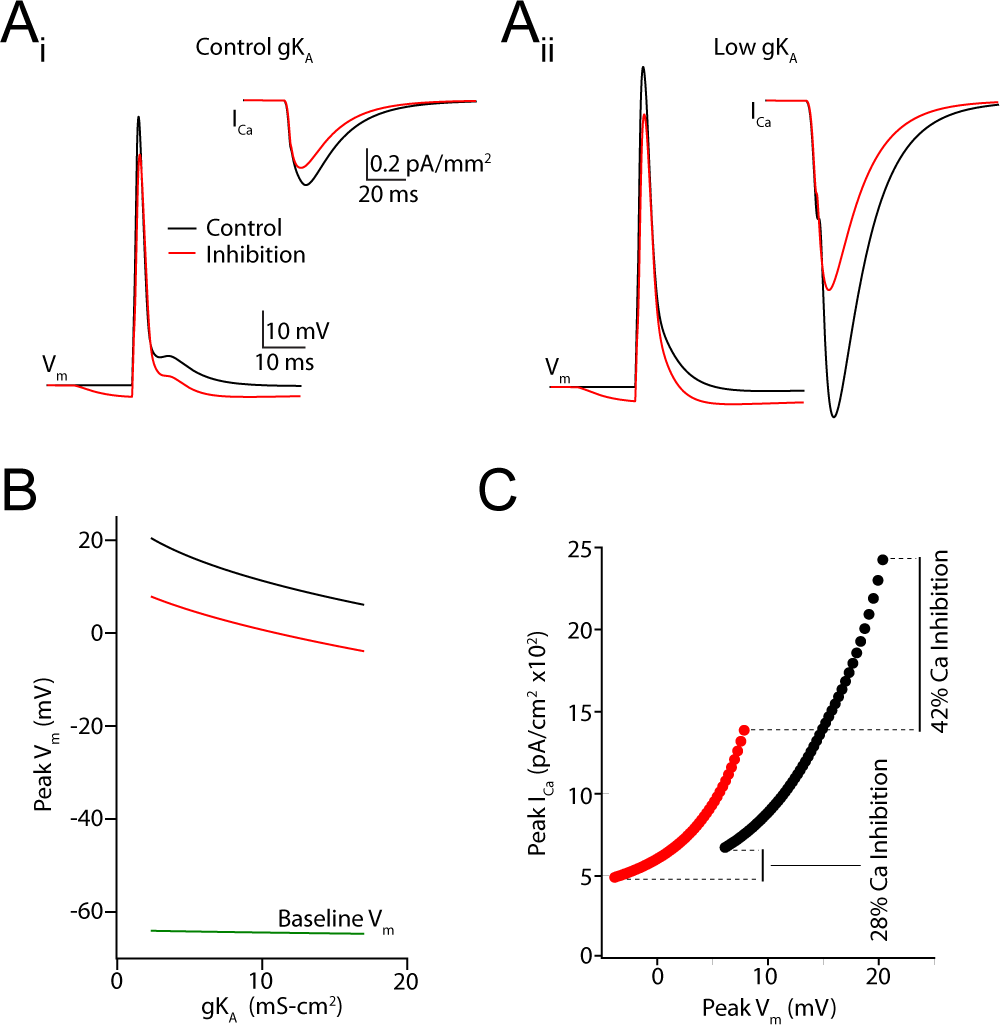
Computational simulations reveal mechanisms underlying potassium channel-dependent regulation of inhibition. (**A**) Simulated action potential waveforms and Ca2+ currents under control conditions (black) and when preceded 15 ms by a GABAergic IPSP for a control value of gK_A_ (A1) and lowered gK_A_ (A2). (**B**) Relationship between peak membrane potential during the AP and the magnitude of gK_A_ for control conditions (black) and following GABAergic inhibition (red). The baseline resting membrane potential is shown in green. (C) Relationship between peak Ca2+ current and peak membrane potential during the AP for control conditions (black) and following GABAergic inhibition (red). Lines traverse varying magnitude of gK_A_ values.

## Discussion

Voltage-gated potassium channels are widely recognized as key modulators of neuronal excitability as well as synaptic integration and plasticity (22, 24, 26, 32–34). In the present work, we have described a role for these channels in the regulation of GABAergic control over dendritic Ca2+ signaling. Using a combination of electrophysiology, 2-photon Ca2+ imaging, and focal GABA uncaging, we show that blocking potassium channels either pharmacologically or via activity-dependent inactivation enhances both bAP-evoked Ca2+ influx and GABAergic inhibition of these transients in the apical dendrites of L5PNs. Our results demonstrate that dendritic inhibition is highly regulated by the expression of voltage-dependent channels near the site of synaptic input.

Dendritic potassium channels comprise a diverse molecular group, including both Kv1-, Kv3, and Kv4-type channels (26, 35). The observation that brief trains of APs appear to rapidly inactivate the channels regulating dendritic inhibition suggests a contribution from A-type conductances, known to be highly sensitive to both membrane depolarization and 4-AP (30, 36–38). Several previous studies have implicated A-type channels in the regulation of both dendritic excitability and glutamatergic synaptic integration. In both CA1 and cortical pyramidal neurons, the presence of A-type channels limits the spread of voltage between distinct compartments, such as the distal and proximal apical dendrite, regulating both the back propagation of action potentials and the spread of synaptically evoked dendritic spikes (23, 24, 31, 34, 39, 40). Our study suggests that these channels similarly restrict the efficacy of GABAergic inhibition, enabling A-type channels to serve as dendritic “shock absorbers”, limiting the impact of synaptic inputs from all sources (41). An intriguing possibility is that voltage-gated potassium channels asymmetrically regulate excitation and inhibition, potentially leading to moment-to-moment alterations of the balance between these opposing drives, a hypothesis whose examination will require additional studies.

In addition to regulating dendritic excitability, voltage-dependent potassium channels have been implicated in shaping long-term plasticity of glutamatergic synapses. In particular, studies have focused on spike-timing dependent plasticity (STDP), where bAPs can potentiate or depress synaptic inputs depending on their relative timing to synaptic activity (42, 43). For example, EPSPs in CA1 pyramidal neuron dendrites can inactivate A-type channels, enhancing the dendritic invasion of somatic action potentials and subsequent plasticity (22). Recent experimental and computational studies have also suggested a key role for GABAergic inhibition in spike-timing dependent plasticity (44–47). For example, focal activation of GABAergic synapses in CA1 dendrites was shown to convert long-term potentiation to depression due to negative regulation of dendritic Ca2+ influx (44). Together, these various findings suggest that the interaction of NMDARs, A-type channels, and GABAergic inhibition may strongly contribute to the development and maintenance of cortical circuits.

It is intriguing to speculate that expression patterns of potassium channels may also explain some of the recent diversity in studies examining GABAergic control of dendritic ΔCa2+. Previous work from our lab showed that inhibition could be highly compartmentalized in layer 2/3 pyramidal neurons, with neighboring spines exhibiting markedly different amounts of inhibition (10). In contrast, work from other groups has shown that more broad dendritic inhibition can occur in L5PNs and hippocampal CA1 pyramidal neurons (13–15). Differential expression and recruitment of voltage-gated conductances, such as KA channels, would be expected to contribute to the heterogeneity of inhibitory function across cell types.

Our computational modeling provides additional insight into the biophysical mechanism underlying the interaction of potassium channels and GABAergic inhibition of Ca2+ influx. We found that decreasing gK_A_ increased the peak depolarization of the AP, producing a supralinear increase in Ca2+ current and a subsequent enhancement of Ca2+ inhibition. The relationship between action potential waveform and calcium influx has been demonstrated previously in presynaptic terminals (48, 49). Our results highlight a similar phenomenon in dendrites and indicate that a major contributor to the potency of GABAergic influence over dendritic Ca2+ signaling is the relationship between bAP waveform and Ca2+ channel activation. Our results differ from previously published work in hippocampal PNs that found increased Ca2+ inhibition for smaller Ca2+ transients (14). In contrast, we find that in both experimental and simulation data, GABAergic inhibition is more potent when the Ca2+ transient is larger following reduction of potassium channel conductance. The disparate findings likely reflect a complicated relationship between total membrane conductance, depolarization, and Ca2+ channel activation that may vary between cell types and experimental conditions.

Finally, we found that the voltage-dependent inactivation of potassium channels allows for the enhancement of GABAergic inhibition in the presence of high frequency somatic spike generation. This suggests that dendritic inhibition may exert greater control over Ca2+ signaling during periods of high network activity or somatic depolarization, essentially acting as a source of homeostatic control. These findings are consistent with previous experimental and computational studies demonstrating the activity-dependent amplification of trains of bAPs in L5PN apical dendrites (31, 50, 51). Our results suggest that the dynamic properties of active dendritic conductances enable the alteration of GABAergic inhibition over short millisecond time frames, providing the basis for a context-dependent, flexible role of GABAergic signaling in shaping biochemical signaling in dendrites.

## Materials and Methods

### Slice Preparation

All animal handling was performed in accordance with guidelines approved by the Yale Institutional Animal Care and Use Committee and federal guidelines. For GABA uncaging experiments, subjects were male wild-type C57-BL6 mice, ages P30-40 (Harlan). For optogenetic experiments, subjects were male and female SOM-Cre mice, ages P30-40 (IMSR Cat# JAX:013044, RRID:IMSR_JAX:013044). Under isofluorane anesthesia, mice were decapitated and coronal slices (300 μm thick) containing primary visual cortex were cut in ice cold external solution containing (in mM): 110 choline, 25 NaHCO3, 1.25 NaH2PO4, 2.5 KCl, 7 MgCl2, 0.5 CaCl2, 20 glucose, 11.6 sodium ascorbate, and 3.1 sodium pyruvate, bubbled with 95% O2 and 5% CO2. After an incubation period of 20 minutes at 34°C, slices were transferred to artificial cerebrospinal fluid (ACSF) containing in (mM): 127 NaCl, 25 NaHCO3, 1.25 NaH2PO4, 2.5 KCl, 1 MgCl2, 2 CaCl2, and 20 glucose bubbled with 95% O2 and 5% CO2 and maintained at room temperature (20-22°C) for at least 20 min until use.

### Electrophysiology and imaging

Experiments were conducted at room temperature in a submersion type recording chamber. Whole-cell patch clamp recordings were obtained from layer 5 pyramidal neurons (500 μm to 600 μm from the pial surface) identified with video infrared-differential interference contrast. For current-clamp recordings, glass electrodes (2-4 ΜΩ tip resistance) were filled with internal solution containing (in mM): 135 KMeSO3, 10 HEPES, 4 MGCl2, 4 Na2ATP, 0.5 NaGTP, and 10 sodium creatine phosphate, adjusted to pH 7.3 with KOH. For Ca2+ imaging experiments, red fluorescent Alexa Fluor-568 (40 μM) and green fluorescent Ca2+-sensitive Fluo-5F (300 μM) were included in the pipette solution to visualize cell morphology and changes of intracellular Ca2+ concentration, respectively. Electrophysiological recordings were made using a Multiclamp 700B amplifier (Molecular Devices), filtered at 4 kHz, and digitized at 10 kHz. For all recordings, membrane potential was adjusted to -64 mV using current injection through the pipette.

Two-photon imaging was performed with a custom-modified Olympus BX51-WI microscope, including components manufactured by Mike's Machine Company. Fluorophores were excited using 840 nm light from a pulsed titanium-sapphire laser. Emissions were separated using appropriate optical filters (Chroma, Semrock) and collected by photomultiplier tubes (Hamamatsu). A mechanical shutter was placed in front of the collectors to prevent damage during blue light stimulation. For Ca2+ imaging, signals were collected during a 500 Hz line scan across a spine and neighboring dendritic shaft on the main apical trunk 100 μΔι to 150 μm from the cell body. Back-propagating action potentials (bAPs) were evoked using a brief depolarizing current pulse (0.5 ms, 1.5-2.5 nA) through the recording pipette. Trials including bAP alone, IPSP-bAP, and IPSP alone were interleaved with a 45 second inter-trial interval. In a subset of experiments, trains of action potentials at 50 Hz and 100 Hz were elicited by current pulse injections through the recording pipette, ending 20 ms prior to a single current pulse. In this case, trials including single bAP alone, train-bAP alone, IPSP-bAP, train-IPSP-bAP, IPSP alone, train-IPSP, and train alone were interleaved with a 45 ms inter-trial interval. Fluorescent traces were computed for individual cells as the average of 10 trials.

Reference frame scans were taken between each acquisition to correct for small spatial drift over time. Ca2+ signals were first quantified as changes in green fluorescence from baseline normalized to the average red fluorescence ΔG/R). To permit comparison of the imaging data across various microscope configurations, we expressed fluorescence changes as the fraction of the G/R ratio measured in saturating Ca2+ ΔG/Gsat).

### Data acquisition and analysis

Imaging and physiology data were acquired using custom software written in MATLAB. Off-line analysis was performed using custom routines written in MATLAB (MATLAB, RRID:SCR_001622) and IgorPro (Wavemetrics Software, RRID:SCR_000325). Ca2+ responses were calculated as the integral of the fluorescence transient over the first 100 ms after bAP initiation. In order to enable comparisons across cells, Ca2+ inhibition was expressed as in previous studies (10) as ΔCa2+CtlΔCa2+Inh)MCa2+al· All statistical comparisons were made using the non-parametric Wilcoxon matched pairs signed rank test in GraphPad Prism version 7.01 (GraphPad Prism, RRID:SCR_002798) unless otherwise noted.

### Pharmacology

For all GABA uncaging experiments, ACSF included 3 μM CGP-55845 hydrochloride (Tocris Cat. No. 1248) to block GABAB receptors, 10 μM (R)-CPP (Tocris Cat. No. 0247) to block NMDA receptors, and 10 μM NBQX disodium salt (Tocris Cat. No. 1044) to block AMPA receptors. For a subset of experiments, the ACSF included 5 mM 4-aminopyradine (Tocris Cat. No. 0940) or 100 μM picrotoxin (Tocris Cat. No 1128). Local application of 25 mM 4-AP was achieved using a glass puffer pipette (< 2 μm tip) coupled to a Picospritzer. Drugs were ejected continuously with 10-17 psi, and pipettes were position 30-70 μm from the targeted structure at the surface of the slice. In experiments where one-photon uncaging was performed with local drug application, 10.8 μM RuBi-GABA was included in the puffer pipette. In a subset of cells, somatic current injections elicited bursts of action potentials in the presence of 4-AP and were excluded from subsequent analysis.

Visible light-evoked GABA uncaging was accomplished using RuBi-GABA (10.8 μM) bath-applied in the ACSF (29). We overfilled the back aperture of the microscope objective (60x, 1.0 NA) with collimated blue light from a fiber-coupled 473 nm laser. Spherical aberrations due to fiber-coupling resulted in a 15-20 μm diameter disc of light at the focal plane centered on the field of view. A brief (0.5 ms) pulse of light (1-2 mW at the sample) reliably evoked uncaging-evoked IPSPs. For Ca2+ imaging experiments, a blue light photo-artifact was corrected by subtracting fluorescence traces on uncaging-alone trials from those with Ca2+ imaging. For all experiments, GABA uncaging occurred 15 ms prior to bAP initiation.

### ChR2 Expression and Activation

To stimulate SOM-INs, SOM-Cre mice were injected 13-23 days prior to slice preparation into the primary visual cortex with recombinant adeno-associated virus (AAV) driving conditional expression of a ChR2-eYFP fusion protein under the Ef1a-promoter (AAV-DIO-Ef1a-ChR2-EYFP)(UNC Vector Core). Optogenetic stimulation was accomplished using the same light source and path as one-photon GABA uncaging (see above). Brief (2-3 mW, 0.5 ms) pulses were used to stimulate SOM-INs 15 ms prior to bAP initiation.

### NEURON Modeling

Multi-compartment time-dependent simulations were run using NEURON v7.4 (NEURON, RRID:SCR_005393, available free at http://neuron.med.yale.edu) and analyzed using custom scripts written in Jupyter Notebooks 4.1.0 using Python 3.5.2. We modified a previously published ball and stick model adding two apical dendrites and dividing the main apical dendrite into 100 segments (length 5 □m each) (10). Sodium channels (4 mS/ cm2) and Hodgkin-Huxley style potassium channels (0.1 mS/cm2) were constant throughout the dendrite. A single dendritic spine (1 □m diameter) was attached to the apical dendrite 122.5 □m from the cell body by a neck (1 □m length, 0.07 □m diameter). A GABAergic synapse (utilizing GABAA receptors) was modeled as an exponential synapse with the NEURON Exp2Syn mechanism (Gmax=2 nS, τ1=5 ms, τ2 =74 ms), contacting the dendritic shaft located 122.5 μm from the cell body. Chloride reversal potential was set to -70 mV. We modeled an A-type potassium conductance using a previously published channel definition that fits observed currents in distal dendrites (52, 53). The A-type channel densities were set at 0 at the soma and proximal dendritic segment and increased linearly with distance. Maximum conductance in the distal segment of the dendrite was varied from 10 mS/cm2 to 70 mS/cm2. A previously published medium voltage-gated calcium channel was inserted into the dendritic spine and neighboring dendrite (1e-7 mS/cm) such that currents through these channels would minimally impact membrane potential (54). In order to reproduce our experimental conditions, an iterative search was conducted to find a somatic current injection that maintained the somatic resting potential at 64.00 +/− 0.001 mV at the cell body for each condition tested. Back-propagating action potentials were generated by current injection to the somatic compartment, and inhibitory conductance preceded action potentials by 15 ms. Similar to our experiments, we quantified calcium flux over a 100 ms window in order to calculate percent calcium change due to inhibition. Data were generated for fixed time steps (implicit Euler, dt= 0.005 ms). To speed up simulation time, simulations were run in parallel using the built-in message passing interface of NEURON.

## Acknowledgements

The authors thank Drs. Jess Cardin, Susumu Tomita, and Michael Crair as well as members of the Higley lab for constructive comments during the preparation of this manuscript. Experiments were supported by funding from the March of Dimes (Basil O'Connor Award) and the NIH (R01 MH099045, T32 EY22312).

## Figure Legends

**Supplemental Figure 1.**
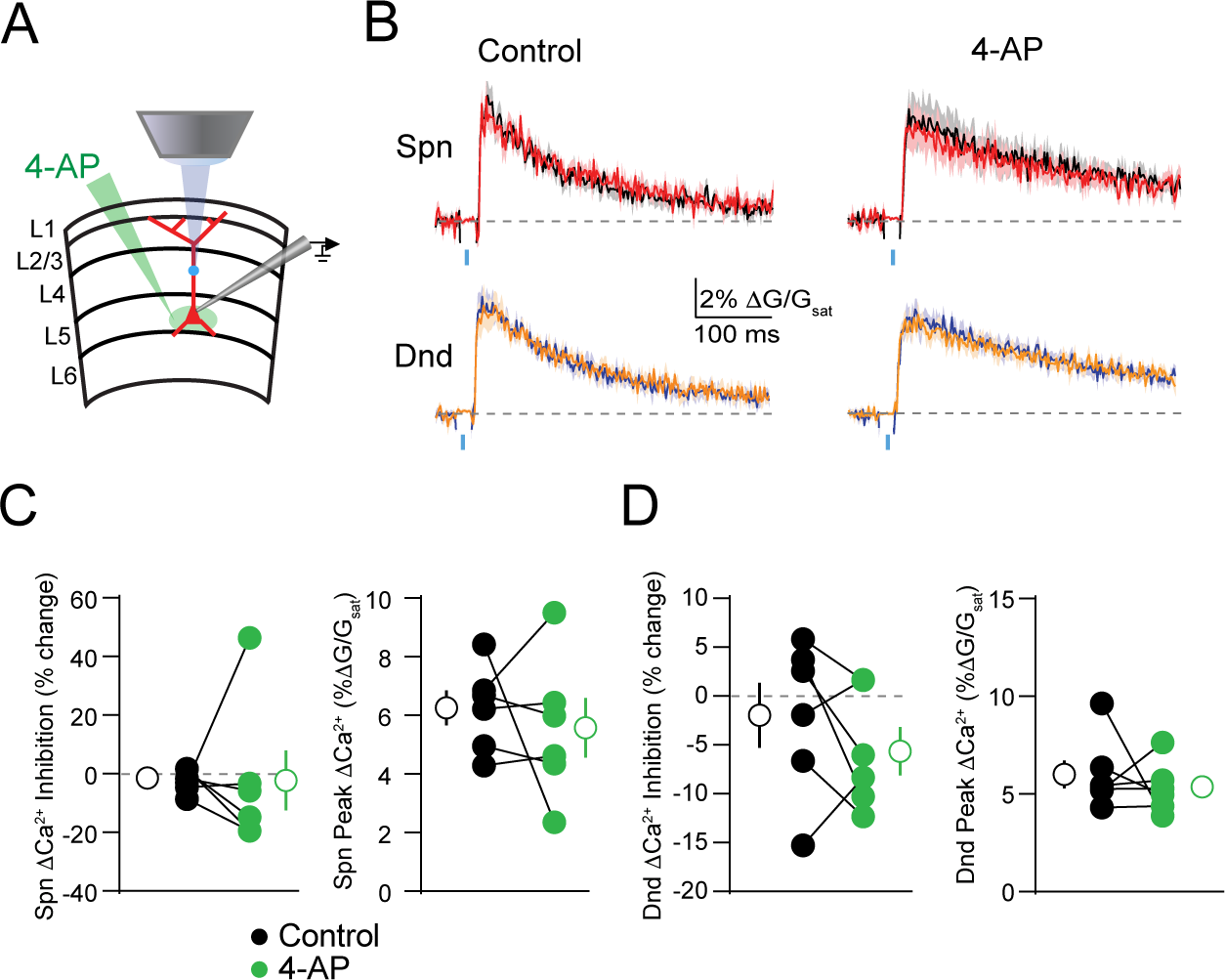
GABAergic inhibition of Δ Ca2+ in the proximal apical dendrite is not affected by somatic potassium channel blockade. (**A**) Schematic of recording and imaging configuration, illustrating somatic location of puffed 4-AP. (**B**) Average ± SEM (n=6) Ca2+ transients for bAP alone or paired with GABA uncaging before (left) and after (right) application of 4-AP to the soma (colors as in Fig. 1). (**C-D**) Population data (n=6) showing the magnitude of Ca2+ inhibition and peak bAP-evoked Ca2+ for spines (C) and neighboring dendritic shafts (D) in control (black) and 4-AP (green) conditions (Wilcoxon matched-pairs signed rank test, p>0.05).

**Supplemental Figure 2.**
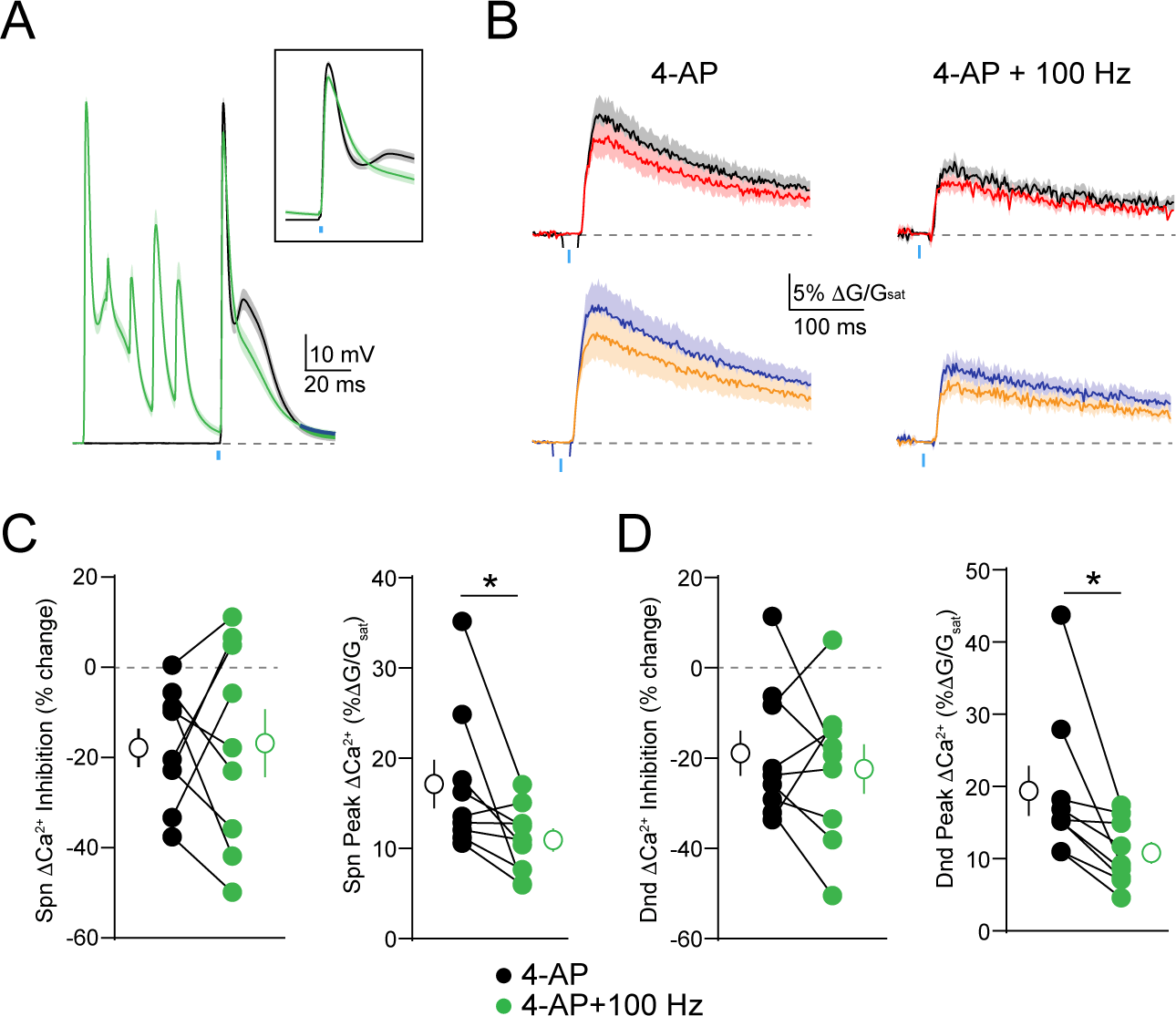
Activity-dependent enhancement of GABAergic inhibition is occluded by blockade of potassium channels. (**A**) Average ± SEM (n=10) of somatic recordings for a single action potential (black) or when preceded 20 ms by a 100 Hz train of action potentials (green), in the presence of 5 mM 4-AP. Inset shows change in spike waveform. (**B**) Average ± SEM (n=10) Ca2+ transients for bAP alone or paired with GABA uncaging, presented either singly (left) or following a 100 Hz train of action potentials (right), in the presence of 4-AP (colors as in Fig. 1). (**C-D**) Population data (n=10) showing the magnitude of Ca2+ inhibition and peak bAP-evoked Ca2+ for spines (C) and neighboring dendritic shafts (D), presented either singly (black) or following a 100 Hz train of action potentials (green), in the presence of 4-AP (Wilcoxon matched-pairs signed rank test, *p<0.05).

## References

1. Zhu Y, Stornetta RL, & Zhu JJ (2004) Chandelier cells control excessive cortical excitation: characteristics of whisker-evoked synaptic responses of layer 2/3 nonpyramidal and pyramidal neurons. The Journal of neuroscience : the official journal of the Society for Neuroscience 24(22):5101–5108.

2. Higley MJ & Contreras D (2006) Balanced excitation and inhibition determine spike timing during frequency adaptation. The Journal of neuroscience : the official journal of the Society for Neuroscience 26(2):448–457.

3. Pouille F & Scanziani M (2001) Enforcement of temporal fidelity in pyramidal cells by somatic feedforward inhibition. Science 293(5532):1159–1163.

4. Wehr M & Zador AM (2003) Balanced inhibition underlies tuning and sharpens spike timing in auditory cortex. Nature 426(6965):442–446.

5. Beaulieu C & Somogyi P (1990) Targets and Quantitative Distribution of GABAergic Synapses in the Visual Cortex of the Cat. The European journal of neuroscience 2(10):896.

6. Miles R, Toth K, Gulyas AI, Hajos N, & Freund TF (1996) Differences between Somatic and Dendritic Inhibition in the Hippocampus. Neuron 16(4):815–823.

7. Lovett-Barron M, et al. (2012) Regulation of neuronal input transformations by tunable dendritic inhibition. Nature neuroscience 15(3):423–430, S421-423.

8. Bloss EB, et al. (2016) Structured Dendritic Inhibition Supports Branch-Selective Integration in CA1 Pyramidal Cells. Neuron 89(5):1016–1030.

9. Tsubokawa H & Ross WN (1996) IPSPs modulate spike backpropagation and associated [Ca2+]i changes in the dendrites of hippocampal CA1 pyramidal neurons. Journal of neurophysiology 76(5):2896–2906.

10. Chiu CQ, et al. (2013) Compartmentalization of GABAergic inhibition by dendritic spines. Science 340(6133):759–762.

11. Palmer L, Murayama M, & Larkum M (2012) Inhibitory Regulation of Dendritic Activity in vivo. Frontiers in neural circuits 6:26.

12. Kanemoto Y, et al. (2011) Spatial distributions of GABA receptors and local inhibition of Ca2+ transients studied with GABA uncaging in the dendrites of CA1 pyramidal neurons. PloS one 6(7):e22652.

13. Marlin JJ & Carter AG (2014) GABA-A receptor inhibition of local calcium signaling in spines and dendrites. The Journal of neuroscience : the official journal of the Society for Neuroscience 34(48):15898–15911.

14. Mullner FE, Wierenga CJ, & Bonhoeffer T (2015) Precision of Inhibition: Dendritic Inhibition by Individual GABAergic Synapses on Hippocampal Pyramidal Cells Is Confined in Space and Time. Neuron 87(3):576–589.

15. Stokes CC, Teeter CM, & Isaacson JS (2014) Single dendrite-targeting interneurons generate branch-specific inhibition. Frontiers in neural circuits 8:139.

16. Cook EP & Johnston D (1999) Voltage-dependent properties of dendrites that eliminate location-dependent variability of synaptic input. Journal of neurophysiology 81(2):535–543.

17. Johnston D & Narayanan R (2008) Active dendrites: colorful wings of the mysterious butterflies. Trends in neurosciences 31(6):309–316.

18. Miller JP, Rall W, & Rinzel J (1985) Synaptic amplification by active membrane in dendritic spines. Brain research 325(1–2):325–330.

19. Poirazi P & Mel BW (2001) Impact of active dendrites and structural plasticity on the memory capacity of neural tissue. Neuron 29(3):779–796.

20. Shepherd GM, et al. (1985) Signal enhancement in distal cortical dendrites by means of interactions between active dendritic spines. Proceedings of the National Academy of Sciences of the United States of America 82(7):2192–2195.

21. Gasparini S (2011) Distance- and activity-dependent modulation of spike back-propagation in layer V pyramidal neurons of the medial entorhinal cortex. Journal of neurophysiology 105(3):1372–1379.

22. Hoffman DA, Magee JC, Colbert CM, & Johnston D (1997) K+ channel regulation of signal propagation in dendrites of hippocampal pyramidal neurons. Nature 387(6636):869–875.

23. Cai X, et al. (2004) Unique roles of SK and Kv4.2 potassium channels in dendritic integration. Neuron 44(2):351–364.

24. Harnett MT, Xu NL, Magee JC, & Williams SR (2013) Potassium channels control the interaction between active dendritic integration compartments in layer 5 cortical pyramidal neurons. Neuron 79(3):516–529.

25. Ramakers GM & Storm JF (2002) A postsynaptic transient K(+) current modulated by arachidonic acid regulates synaptic integration and threshold for LTP induction in hippocampal pyramidal cells. Proceedings of the National Academy of Sciences of the United States of America 99(15):10144–10149.

26. Carrasquillo Y, Burkhalter A, & Nerbonne JM (2012) A-type K+ channels encoded by Kv4.2, Kv4.3 and Kv1.4 differentially regulate intrinsic excitability of cortical pyramidal neurons. The Journal of physiology 590(16):3877–3890.

27. Burkhalter A, Gonchar Y, Mellor RL, & Nerbonne JM (2006) Differential expression of I(A) channel subunits Kv4.2 and Kv4.3 in mouse visual cortical neurons and synapses. The Journal of neuroscience : the official journal of the Society for Neuroscience 26(47):12274–12282.

28. Jinno S, Jeromin A, & Kosaka T (2005) Postsynaptic and extrasynaptic localization of Kv4.2 channels in the mouse hippocampal region, with special reference to targeted clustering at gabaergic synapses. Neuroscience 134(2):483–494.

29. Rial Verde EM, Zayat L, Etchenique R, & Yuste R (2008) Photorelease of GABA with Visible Light Using an Inorganic Caging Group. Frontiers in neural circuits 2:2.

30. Bekkers JM (2000) Distribution and activation of voltage-gated potassium channels in cell-attached and outside-out patches from large layer 5 cortical pyramidal neurons of the rat. The Journal of physiology 525 Pt 3:611–620.

31. Kim J, Wei DS, & Hoffman DA (2005) Kv4 potassium channel subunits control action potential repolarization and frequency-dependent broadening in rat hippocampal CA1 pyramidal neurones. The Journal of physiology 569(Pt 1):41–57.

32. Foeger NC, Norris AJ, Wren LM, & Nerbonne JM (2012) Augmentation of Kv4.2-encoded currents by accessory dipeptidyl peptidase 6 and 10 subunits reflects selective cell surface Kv4.2 protein stabilization. The Journal of biological chemistry 287(12):9640–9650.

33. Magee JC & Carruth M (1999) Dendritic voltage-gated ion channels regulate the action potential firing mode of hippocampal CA1 pyramidal neurons. Journal of neurophysiology 82(4):1895–1901.

34. Losonczy A, Makara JK, & Magee JC (2008) Compartmentalized dendritic plasticity and input feature storage in neurons. Nature 452(7186):436–441.

35. Serodio P, Kentros C, & Rudy B (1994) Identification of molecular components of A-type channels activating at subthreshold potentials. Journal of neurophysiology 72(4):1516–1529.

36. Serodio P & Rudy B (1998) Differential expression of Kv4 K+ channel subunits mediating subthreshold transient K+ (A-type) currents in rat brain. Journal of neurophysiology 79(2):1081–1091.

37. Clark BD, et al. (2008) DPP6 Localization in Brain Supports Function as a Kv4 Channel Associated Protein. Frontiers in molecular neuroscience 1:8.

38. Korngreen A & Sakmann B (2000) Voltage-gated K+ channels in layer 5 neocortical pyramidal neurones from young rats: subtypes and gradients. The Journal of physiology 525 Pt 3:621–639.

39. Kim J, Jung SC, Clemens AM, Petralia RS, & Hoffman DA (2007) Regulation of dendritic excitability by activity-dependent trafficking of the A-type K+ channel subunit Kv4.2 in hippocampal neurons. Neuron 54(6):933–947.

40. Frick A, Magee J, Koester HJ, Migliore M, & Johnston D (2003) Normalization of Ca2+ signals by small oblique dendrites of CA1 pyramidal neurons. The Journal of neuroscience : the official journal of the Society for Neuroscience 23(8):3243–3250.

41. Yuste R (1997) Potassium channels. Dendritic shock absorbers. Nature 387(6636):851–853.

42. Magee JC & Johnston D (1997) A synaptically controlled, associative signal for Hebbian plasticity in hippocampal neurons. Science 275(5297):209–213.

43. Markram H, Lubke J, Frotscher M, & Sakmann B (1997) Regulation of synaptic efficacy by coincidence of postsynaptic APs and EPSPs. Science 275(5297):213–215.

44. Hayama T, et al. (2013) GABA promotes the competitive selection of dendritic spines by controlling local Ca2+ signaling. Nature neuroscience 16(10):1409–1416.

45. Wilmes KA, Sprekeler H, & Schreiber S (2016) Inhibition as a Binary Switch for Excitatory Plasticity in Pyramidal Neurons. PLoS computational biology 12(3):e1004768.

46. Cichon J & Gan WB (2015) Branch-specific dendritic Ca(2+) spikes cause persistent synaptic plasticity. Nature 520(7546):180–185.

47. Paille V, et al. (2013) GABAergic circuits control spike-timing-dependent plasticity. The Journal of neuroscience : the official journal of the Society for Neuroscience 33(22):9353–9363.

48. Zucker RS, Delaney KR, Mulkey R, & Tank DW (1991) Presynaptic calcium in transmitter release and posttetanic potentiation. Annals of the New York Academy of Sciences 635:191–207.

49. Augustine GJ (1990) Regulation of transmitter release at the squid giant synapse by presynaptic delayed rectifier potassium current. The Journal of physiology 431:343–364.

50. Larkum ME, Zhu JJ, & Sakmann B (1999) A new cellular mechanism for coupling inputs arriving at different cortical layers. Nature 398(6725):338–341.

51. Grewe BF, Bonnan A, & Frick A (2010) Back-Propagation of Physiological Action Potential Output in Dendrites of Slender-Tufted L5A Pyramidal Neurons. Frontiers in cellular neuroscience 4:13.

52. Migliore M, Hoffman DA, Magee JC, & Johnston D (1999) Role of an A-type K+ conductance in the back-propagation of action potentials in the dendrites of hippocampal pyramidal neurons. Journal of computational neuroscience 7(1):5–15.

53. Acker CD & Antic SD (2009) Quantitative assessment of the distributions of membrane conductances involved in action potential backpropagation along basal dendrites. Journal of neurophysiology 101(3): 1524–1541.

54. Polsky A, Mel BW, & Schiller J (2004) Computational subunits in thin dendrites of pyramidal cells. Nature neuroscience 7(6):621–627.

